# Genome evolution during the domestication of an antibiotic-producing *Streptomyces* strain

**DOI:** 10.64898/2026.08.28.747838

**Authors:** John T. Munnoch, Daniel E. Larcombe, Rebecca E. McHugh, John Bruce, Kirsty Robb, James T. Croxford, Angelika B. Kiepas, Juan Pablo Gomez-Escribano, Nicola A. Crowhurst, Andrew J. Collis, Steven G. Kendrew, Benjamin D. Huckle, Barrie Wilkinson, Iain S. Hunter, Paul A. Hoskisson

## Abstract

The domestication of *Streptomyces* species for antibiotic production involves long-term, iterative mutagenesis and selection, yet the genomic changes driving enhanced production remain unclear. Analysis of five strains from an industrial lineage of *Streptomyces clavuligerus* using comparative genomics, transcriptomics and phenotypic profiling for dynamic genome architectures with plasmid integration events and chromosomal rearrangements, alongside the accumulation of mutations affecting metabolic pathways and global gene regulation. These changes increased precursor supply and reprogrammed transcription leading to enhanced clavulanic acid production but reduced catabolic flexibility. Complementation experiments confirmed the functional impacts of specific mutations. These findings reveal that artificial selection shapes genome evolution in industrial strains, balancing production gains with metabolic trade-offs. This work will likely inform rational design of *Streptomyces* strains for improved natural product production in industry while highlighting the constraints imposed by domestication on metabolic versatility. More broadly it shows that many of the evolutionary processes in industrial strain improvement programmes mirror those at play during natural selection.

## Introduction

Understanding the adaptation of populations to their environment is a fundamental issue in biology, when introduction of genetic variation within a population is followed by selection. The process of domestication inspired Darwin (Darwin 1859; Gregory 2009) to draw important parallels between the process of adaptation with strong artificial selection by humans and evolution by natural selection. Humans have been unintentionally domesticating microorganisms for thousands of years (Steensels et al. 2019) and, more recently, industrial strain improvement programmes have been selecting microorganisms for improved performance for a range of industrial and biotechnological applications. The development of high-throughput mutagenesis screens for the improvement of many industrial microorganisms accelerated this process but it has largely remained an empirical science.

### Streptomyces

bacteria are the industrial workhorses for production of a wide range of pharmaceutically useful bioactive natural products such as antimicrobials, immunosuppressives, anti-cancer agents and antihelmintics (Chevrette et al. 2019; Hoskisson and Seipke 2020). For over 70 years the development of *Streptomyces* strains for production of bioactive natural products has been achieved via random mutagenesis using chemical mutagens or UV and/or recombination, followed by selection for increased product formation (Baltz 2011). Whilst this approach to strain improvement is successful at deriving high-producing strains, it is often time consuming and can take years to deliver commercially viable results (Nielsen 1997).

The mutational process that drives adaptive evolution in natural situations is analogous to, but accelerated in, the strain improvement processes for industrial microorganisms where these populations suffer bottlenecks due to the strong selection imposed by humans - which can potentially affect future adaptation (Gregory 2009; Cisneros-Mayoral et al. 2022). These iterative high-throughput mutagenesis and screening programmes deliver a wide range of stochastic mutations, resulting in strains with industrially desirable characteristics. Selection for product formation has led to the identification of a range of enhancing mutations such as increased transcription of biosynthetic genes, an increase in the copy number of biosynthetic gene clusters (BGCs), the loss of competing BGC expression and reduced production of contaminating natural products (Yanai et al. 2006; Medema et al. 2011a; Medema et al. 2011b; Paradkar 2013; Fiedurek et al. 2017). Yet the selection of high-producing strains under a highly defined set of conditions may also result in widespread deleterious mutations occurring across the genome. This can result in loss of function in genes not essential in the highly specialised environment of the industrial fermentation. Whilst many high-titre production strains of *Streptomyces* are supremely adapted to the media, culture conditions and processes for which they have been selected, the accumulation of deleterious mutations in these industrial strains may lead to reduced flexibility of these strains and limit future modification of industrial processes.

Clavulanic acid (CA) is a potent ß-lactamase inhibitor produced by *Streptomyces clavuligerus* that potentiates the antibacterial activity of penicillins and cephalosporins against ß-lactamase-producing resistant bacteria (Paradkar 2013). CA is an established clinical molecule and a member of the WHO essential medicines list (https://list.essentialmeds.org/). When co-formulated with amoxicillin (Augmentin/Co-amoxiclav), it is valuable in the global effort to combat antimicrobial resistance. Industrial strains of *S. clavuligerus* have undergone >45 years of strain improvement to enhance production by numerous industrial CA manufacturers using random mutagenesis and recombination, with some strains producing >100-fold higher levels of CA than the wild-type isolate (Rowlands 1984; Ünsaldı et al. 2017; Cho et al. 2019). The time-consuming nature of the classical strain improvement approaches, which often result in genetic instabilities and diminishing returns in terms of production (Gravius et al. 1993; Petkovic et al. 2006; Cho et al. 2019), suggests that greater insight into the genomic wide changes that give rise to increased production will help design rational approaches to strain improvement.

To better understand the domestication of antibiotic-producing *Streptomyces* an extensive analysis of five strains that comprise the early stages of an authentic CA-producing industrial lineage of *S. clavuligerus* was undertaken. Comparative genomics, transcriptomics and phenotypic profiling enabled key insights into the evolution and domestication of this lineage, identifying genomic plasticity in strains, coupled with mutations and transcriptional changes in central metabolic pathways. Complementation of mutations was sufficient to restore catabolic flexibility in some instances and improve CA production. These data will find utility in rational strain design and development programmes for *Streptomyces* going forward in the fight against antimicrobial resistance.

## Results

### The genomes of industrial *Streptomyces clavuligerus* strains are plastic

The industrial lineage of *S. clavuligerus* has undergone more than 45 years of random mutagenesis and selection for improved CA production at Beechams, SmithKline Beecham and subsequently GlaxoSmithKline (GSK). The initial strain of *S. clavuligerus* ATCC 27064 (also deposited as DSM 738 and NRRL 3585; Higgens and Kastner 1971), was purified to a single CA producing colony, which became *S. clavuligerus* SC2 (**Fig. 1**). Chemical mutagenesis (unspecified in the historical Smithkline Beecham literature) was used to generate a branch point in the lineage, leading to *S. clavuligerus* SC3 and *S. clavuligerus* SC4 (**Fig. 1**). The *S. clavuligerus* SC3 branch was not taken any further in the development process and represents a dead end within this industrial lineage. *S. clavuligerus* SC4 subsequently underwent numerous rounds of mutagenesis/selection to increase CA titre, using chemical (unspecified), UV and ionising radiation, prior to the selection of *S. clavuligerus* SC5 (**Fig. 1**). Additional UV mutagenesis of *S. clavuligerus* SC5 gave rise to the strain

**Fig. 1.**
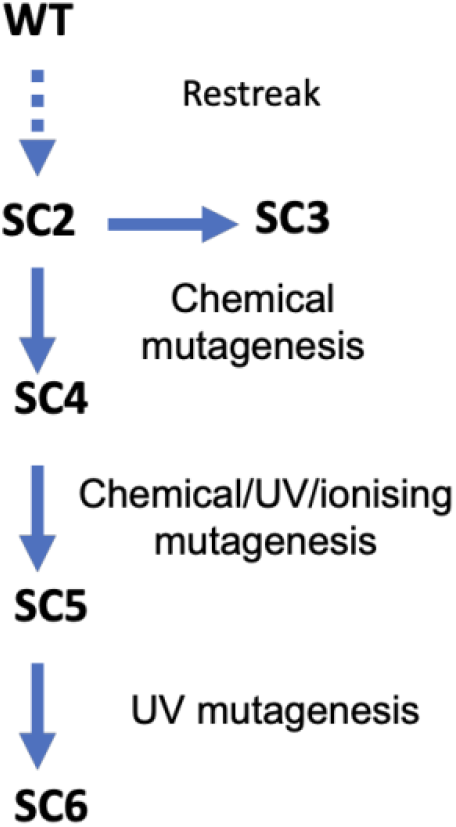
Industrial strain lineage of *Streptomyces clavuligerus*. The strain development route to *S. clavuligerus* SC6 (based on historical data from the GSK archives), along with the mutagenesis methods used to generate the strains used in this study. The strain designations do not represent a single mutagenesis step, but rather a strain that was used industrially for production. Each incremental strain may represent several mutagenesis steps.

### S. clavuligerus

SC6 which was used for the industrial production of CA (**Fig. 1**). Morphologically these strains all undergo sporulation and produce the characteristic green-grey spore pigment associated with this species, with *S. clavuligerus* DSM738 and SC6 strains exhibiting the most consistent sporulation phenotypes (**Fig. 2A**). Selection of strains was based on volumetric CA titre in specialised production media and bioreactors independent of the biomass generated. In laboratory shake flask culture and standard TSB medium, CA yield does not show a steady increase, with later strains performing less well, suggesting that the effect of the selected mutations is dependent on medium and cultivation conditions (**Fig. 2B**).

**Fig. 2.**
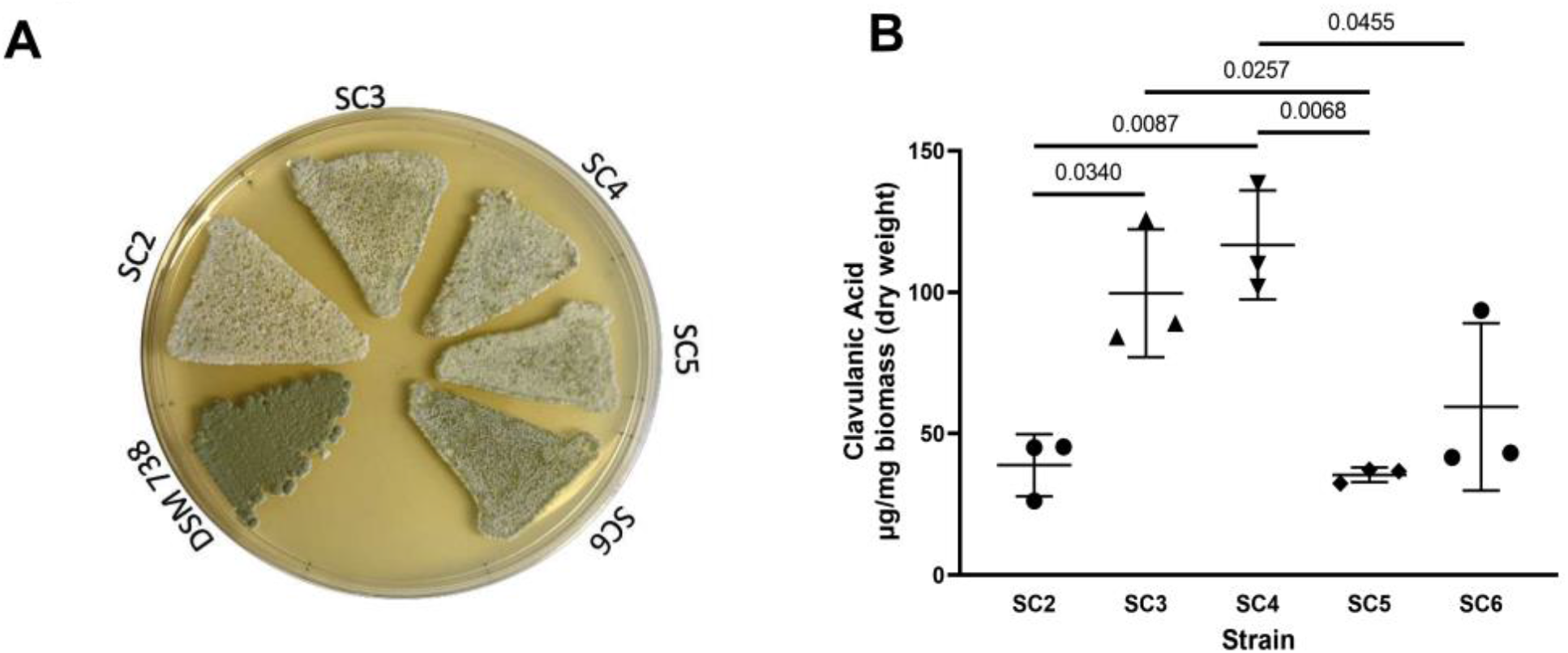
Phenotypic comparison of the *Streptomyces clavuligerus* industrial strain lineage. **A)**. Growth of *S. clavuligerus* DSM 738 (Wild-Type) and the industrial lineage strains on L3M9 agar, showing the changes in spore pigmentation. **B).** Production of clavulanic acid from the industrial strains of *S. clavuligerus* in TSB medium, highlighting the importance of growth medium and process on the evolution and productivity of industrial strains. Means and standard deviations from triplicate experiments are shown. The p-values resulting from two-way ANOVA with Tukey multiple comparison testing are also shown.

To understand the effects of long-term mutagenesis and selection on these strains, whole genome sequencing was undertaken for *S. clavuligerus* SC2 and SC6 using PacBio and Illumina sequencing using a combination of HGAP3 assembly of the PacBio data, followed by hybrid assembly using SPAdes (Bankevich et al., 2012). *S. clavuligerus* SC3, SC4 and SC5 genomes were mapped from Illumina reads using *S. clavuligerus* SC2 as a reference with genomic gaps and rearrangements being confirmed through PCR verification for all strains for genome structural changes. Regions where no sequencing coverage was observed indicating deletions, were also confirmed through PCR, Sanger sequencing and comparison to RNASeq data (**Supp. Fig. 1-10,** and oligonucleotides used in **Supp. Table 1**).

The genomic content of the major *S. clavuligerus* type strains and their plasmids have been sequenced by several groups and were systematically reviewed by (Liras & Martín, 2021) and includes *S. clavuligerus* ATCC 27064 (Broad Institute 2008 - GCA_000154925, Medema et al, [2010] - ADGD01000000, Hwang et al. [2019] – GEO: GSE128216) and Gomez-Escribano et al. (2021) - PRJNA58778 & PRJNA587886, *S. clavuligerus* NRRL 3585 (Wu & Roy 1993) - X54107, Wu et al. 2006)- AY392418 & Song et al. [2010] - ADWJ01000000, *S. clavuligerus* F613-1 Cao et al. [2016] - CP016559 & CP016560, *S. clavuligerus* F1D-5 (GCA_003454 755), *S. clavuligerus* DSM738 (GCF_028752555.1) and *S. clavuligerus* K4567 (GCF_019831915.1, Novartis AG). Reports indicate that the genome comprises an approximate 6.8 Mbp chromosome (Medema et al. 2010; Song et al. 2010) along with a 1.8Mbp megaplasmid (pSCL4; Medema et al. 2010). Three smaller replicons, pSCL1-3, that are between 11 Kbp and 430 Kbp in length, are also found in this species (Wu and Roy 1993; Medema et al. 2010), however they appear to be dynamic in nature with Medema et al., (2010) finding no evidence of pSCL2 and pSCL3 in their assemblies and Hwang et al., (2019) providing no evidence of pSCL1-3. Later sequencing of an isolate derived from the Type strain (DSM 738=ATCC 27064) by Gomez-Escribano et al., (2021), reported the presence of pSCL1-2 and pSCL4 but not pSCL3.

Using the previously published sequences (ADGD01000000; PRJNA58778; PRJNA587886; ADWJ01000000 & GCF_028752555.1) as a guide, mapping and structural analysis of *S. clavuligerus* SC2-SC6 revealed the genome of *S. clavuligerus* genome is plastic, with multiple forms of the genome present in some strains. *S. clavuligerus* SC2 is found in two forms within the population, SC2.1, which largely matches the assembly of Song et al., (2010) with a 6.8 Mbp core genome (6,748,915 bp) with pSCL1 (11,668 bp), pSCL2 (148,877 bp) and pSCL3 (448,731 bp) all assembling as separate replicons (**Fig. 3**). The megaplasmid pSCL4 was found to be substantially smaller in this strain at 835,938 bp rather than 1,797,117 bp, indicating a 961,179 bp loss in content. Additionally, in the assembly approximately half of pSCL1 has recombined onto the left end of pSCL4, while pSCL1 can be detected as an intact replicon (determined from sequencing data and PCR confirmations of these rearrangements **Supp. Fig. 9 and 7** respectively). Structural variants were detected by sequencing and confirmed by PCR for *S. clavuligerus* SC2, this has been denoted SC2.2, where pSCL3 (position 1-325,665 bp) has integrated into the chromosome at position 6,269,822 bp (**Fig. 3**; PCR confirmation of gaps and rearrangements can be found in **Supp. Fig. 1-10**).

**Fig. 3.**
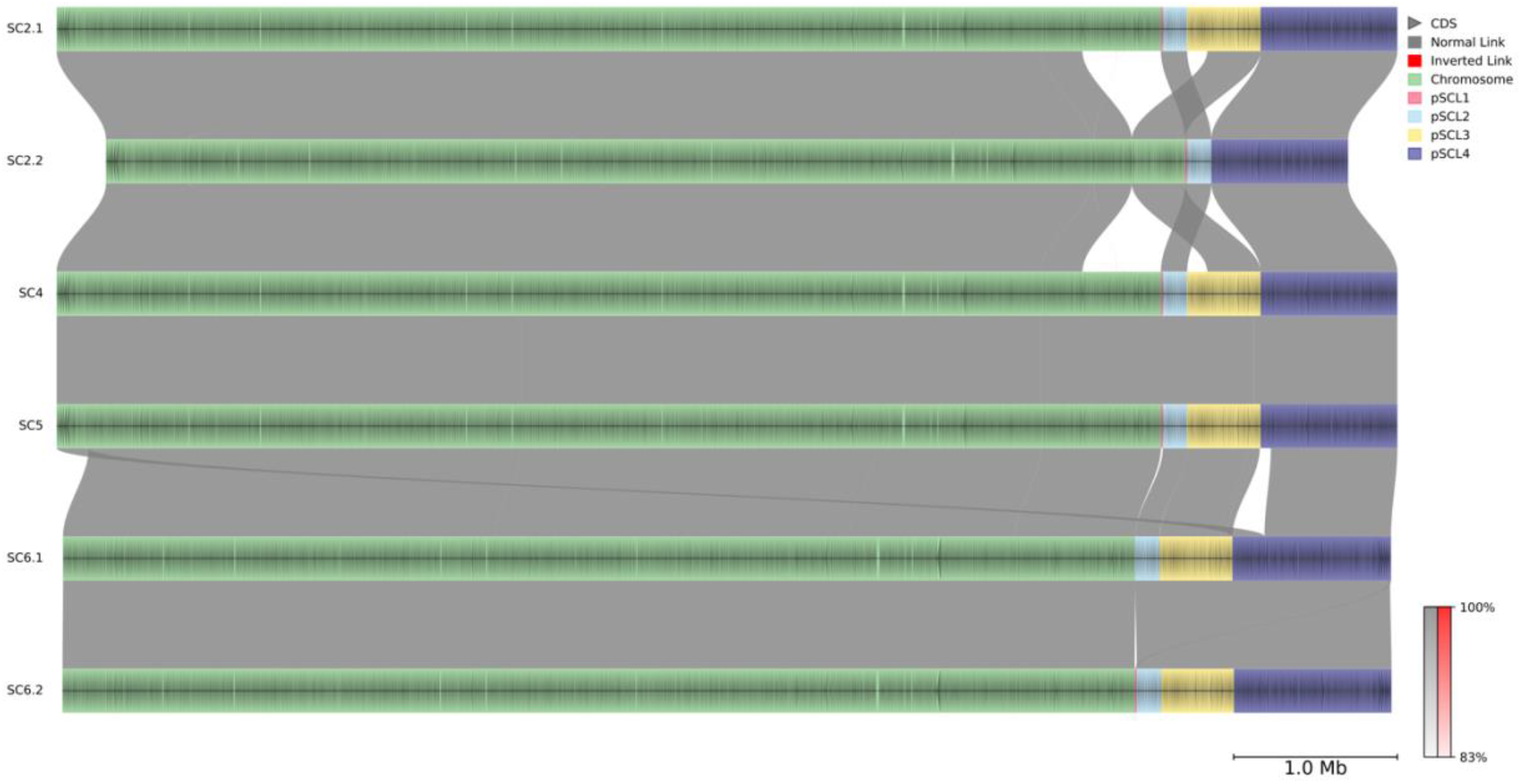
Structural comparison of *Streptomyces clavuligerus* genomes in the industrial lineage. Individual replicons are separated by colours. The chromosome (Green), pSCL1 (Red), pSCL2 (light blue), pSCL3 (yellow), pSCL4 (dark blue). For SC2 and SC6, two versions (.1 and .2) indicate the different structural variants identified through sequencing. Grey colour between the genomes shows synteny in the genomes, with the scale of percentage identity shown on the figure.

The *S. clavuligerus* SC3 strain was selected following chemical mutagenesis of *S. clavuligerus* SC2. The genome sequence of this strain was found to be identical to its progenitor *S. clavuligerus* SC2.1 and analysis of the strain was taken no further as it represents a dead end in the strain lineage.

### S. clavuligerus

SC4 genome has accumulated a total of 15 mutations relative to *S. clavuligerus* SC2, with 10 single nucleotide changes (SNCs) being found in coding sequences, compared to SC2.1 (**Supp. Table 2**). Five of the SNCs result in synonymous changes in coding sequences and five SNCs result in missense mutations: *SCLAV_SC2_00630* (hypothetical protein, putative DNA binding protein) Arg278Trp, *SCLAV_SC2_31875* (Regulatory protein AfsR) Glu121Lys, *SCLAV_SC2_06955* (Heme uptake protein MmpL11) Gly749Asp, *SCLAV_SC2_07245* (hypothetical protein) Val238Ala and *SCLAV_SC2_27080* (Pentalenene oxygenase) Ser135Asn. There are also, four intergenic mutations with no obvious changes to genome features (promoters, terminators etc) and there is a frameshift present in *SCLAV_SC2_18190* (hypothetical protein, putative serine-threonine protein kinase),

The genome architecture of *S. clavuligerus* SC5 is similar *S. clavuligerus* SC2.1 in terms of overall genome structure with a notable difference present in the coverage of pSCL4. There is a region of reduced coverage in pSCL4 (∼42,000 bp to 200,000 bp) with approximately half the sequencing depth, indicating that there are two isoforms of pSCL4 being present in this strain – one with a deletion in the region showing reduced coverage and a version with the replicon intact. All other plasmids are identical to *S. clavuligerus* SC2.1. There are six SNCs and a 48 bp deletion in this strain when compared to *S. clavuligerus* SC4 (**Supp. Table 2**). The intergenic region deleted is at position 2,839,515 nt between two convergently transcribed genes (SCLAV_2401/ SCLAV_SC2_11620, a homologue of *nuoD*, NADH dehydrogenase and SCLAV_2402/ SCLAV_SC2_11625, a putative secreted protein). This deletion which does not appear to affect either CDS, and transcription of both ORFS is evident in the RNA-Seq data and there is no evidence of the presence of stem-loop structures or terminators in deleted region.

In the final strain of the lineage studied to date, *S. clavuligerus* SC6, a distinct form of pSCL4 is found. *S. clavuligerus* SC6.1 (6,553,311 bp) contains the same fragment of pSCL1 observed in SC2.1 transposed on the left end of pSCL4 (1-5,335 bp). In the second form of the genome, *S. clavuligerus* SC6.2, pSCL1 is found as an intact episome (**Supp. Fig. 1-10**). In both forms, the right-hand end of pSCL4 is truncated and the first ∼200kb of the chromosome region is duplicated and transposed on to this end of pSCL4 (PCR confirmations of these rearrangements can be found in **Supp. Fig. 8-10**; A graphical representation of the *S. clavuligerus* SC6 rearrangements can be found in **Supp. Fig. 11**). In *S. clavuligerus* SC6.2, the chromosome is arranged as in *S. clavuligerus* SC6.1, except for pSCL1, where it remains as an episome, suggesting that these genomes are dynamic and plastic in nature with frequent plasmid integration events and rearrangements. Twenty-seven unique SNCs and one deletion were identified in *S. clavuligerus* SC6 that were not present in SC5. All changes are in CDSs, except for one intergenic SNC, found at 1,848,682 nt within the genome. This SNC is located 67 nt upstream of the translational start codon (ATG) of a conserved TetR-like regulator (SCLAV_1553). With respect to the other SNCs, three changes are located in genes encoding putative regulatory proteins (missense mutation in SCLAV_SC2_05705 [Tyr108Cys]; a missense mutation in SCLAV_SC2_07880 [Tyr231Cys]; and missense mutation in SCLAV_SC2_26565 [Val358Gly]). A six bp deletion was found in a gene encoding a putative regulatory protein (SCLAV_SC2_29430). Two missense mutations were also identified in potentially competing natural product biosynthetic gene clusters (BGCs). These were found in two putative polyketide synthases (SCLAV_SC2_00090 [Val3019Ala] and SCLAV_SC2_33825 [Ala365Ser]).

No major deletions were apparent in any of the strains, except for the truncation of pSCL4, rearrangements are present, suggesting that the genomes are dynamic and exist in multiple forms. Extensive changes to overall genome content do not appear to be responsible for the increase in CA production in this lineage, suggesting that CA production changes are brought about through relatively modest genome modifications in the early stages of domestication.

### Industrial *Streptomyces clavuligerus* strains accumulate mutations that reduce catabolic breadth

To analyse the impact of mutations throughout the *S. clavuligerus* lineage, the SNCs across all strains were analysed. It was found that SNCs accumulate with each subsequent step in the lineage from *S. clavuligerus* SC2 and SC6. The greatest number of SNCs was found between *S. clavuligerus* SC5 and SC6. A total of 54 unique SNCs accumulated between the five strains studied (nine of which are synonymous). Gene ontology enrichment analysis (http://geneontology.org/) of these genes indicates that mutations are enriched in genes associated with carbohydrate metabolism, DNA-binding/regulation and secretion (**Supp. Table 3)**.

To analyse the impact of these changes on the physiology of the *S. clavuligerus* strains, Biolog Phenotypic array analysis was undertaken using PM1 Carbon source plates to investigate changes in carbon source utilisation (catabolic breadth) across the lineage. An overall reduction in catabolic capability was found as the *S. clavuligerus* industrial lineage progressed. Thirty carbon sources supported growth of the *S. clavuligerus* strains tested. *S. clavuligeru*s SC2 and *S. clavuligerus* DSM 738 = ATCC27064), were designated as the base line of catabolism (100 %). *S. clavuligerus* SC4, and *S. clavuligerus* SC5 all showed a reduction in catabolic capability compared to the parental strains of *S. clavuligerus* (**Fig. 4A)**. Remarkably, the largest decline in catabolic breadth was observed between *S. clavuligerus* SC4 and SC5, with a subsequent recovery in *S. clavuligerus* SC6, likely due to mutations identified in regulatory genes in *S. clavuligerus* SC6.

**Fig. 4.**
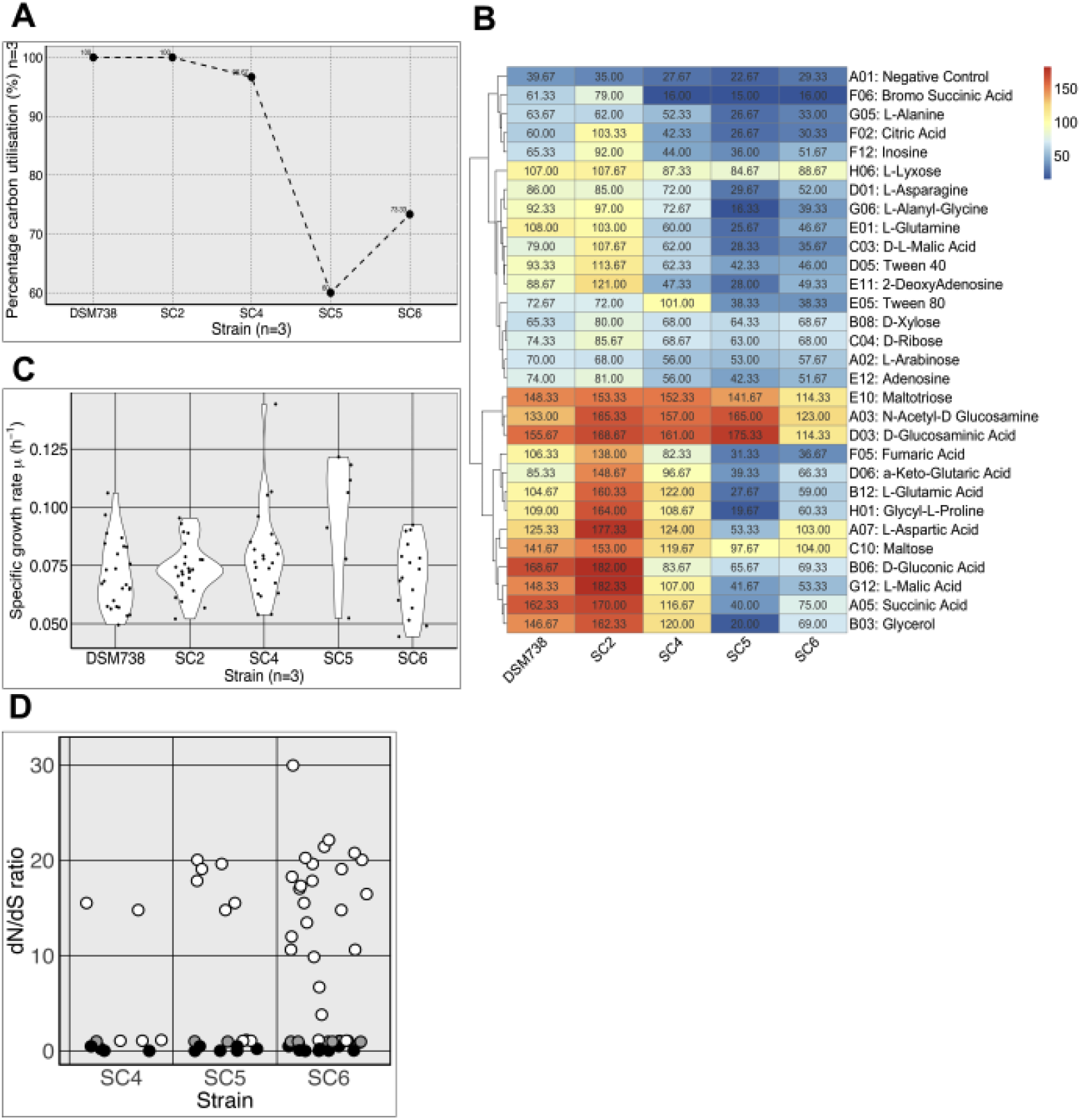
Catabolic function of *Streptomyces clavuligerus* in the industrial lineage. *S. clavuligerus* wild type and strains from the industrial lineage were assessed using Biolog analysis. **A).** Percentage of carbon sources used for *S. clavuligerus* DSM 738 (Wild-Type) and the industrial lineage strains (*S. clavuligerus* SC2-SC6). **B).** Changes in carbon source usage in thirty carbon sources across the industrial lineage. **C).** Violin plots of Specific growth rate (µ, h^-1^) of the *S. clavuligerus* strains across the industrial lineage with 30 different carbon sources. **D).** Selection analysis of mutations in *S. clavuligerus.* dN/dS ratio of genes that have accumulated mutations in the *Streptomyces clavuligerus* in the industrial lineage (SC2 as reference). Black points represent genes with a dN/dS ratio <1 (purifying selection), grey points represent genes with a dN/dS ratio ∼1 (neutral selection), white points represent genes with a dN/dS ratio >1 (positive selection).

The mutation and selection imposed on these strains for certain growth substrates appears to have reduced the breadth of carbon source usage between *S. clavuligerus* SC2 and SC6. The use of starch, maltose and maltotriose in industrial production media is evident when the growth of *S. clavuligerus* SC2 and SC6 is compared (**Fig 4B**). Despite widespread loss of diet breadth, growth capability on N-acetyl-D-glucosamine, D-glucosaminic acid, maltotriose and maltose is retained across the lineage. Recovery of catabolic breadth between *S. clavuligerus* SC5 and SC6 is notable in the recovery of growth on glycerol, L-aspartic acid, L-glutamic acid and glycyl-L-proline. Determination of the specific growth rate (μ) of these strains across 30 carbon sources shows an overall diversification of the growth rate across the permissive carbon sources is likely the result of selection of strains on industrial media containing these or related components (**Fig. 4C**). Whilst reduced catabolic breadth would be considered deleterious in nature, the selection process for these strains by the industrial manufacturer is focused solely on increasing CA titre.

To understand the role of strain selection and how this is reflected in genomic changes in the *S. clavuligerus* industrial lineage, the ratio of amino acid replacing non-synonymous substitutions were compared with synonymous substitutions (dN/dS ratio; *ω*). The dN/dS ratio was calculated for each CDS across the genomes (sequences in **Supp. Table 2**) using Genomegamap (Wilson et al. 2020), revealing strong positive selection (*ω* > 1) in genes with non-synonymous substitutions (**Fig. 4. & Supp. Table 3**). Given this *S. clavuligerus* lineage was developed through artificial selection, focused only on CA titre, selective sweeps likely only purge mutations that negatively affect CA production. This suggests that all extant mutations in the lineage would favour positive selection as strain fitness is being driven based on a single phenotypic trait. Positive selection was seen in genes associated with mycothiol metabolism (*mshA*), beta-oxidation of fatty acids (*fadA*), carbon source transporters (*dasC*), amino acid transporters (*yxeN*) electron transport chain associated diheme c-type cytochrome (*qcrC*) along with several conserved hypothetical protein coding loci. These likely reflect adaptation to industrial fermentation media (containing complex carbon sources and oils from growth substrates such as soya flour) and the oxidative stress conditions found in fermentation processes. Genes with exhibiting synonymous changes (dN/dS<1) included genes associated with DNA repair (*recG*) and nucleotide metabolism (*dut*) suggesting neutral or purifying selection was acting on those genes.

### Selection of strains has altered global transcription in industrial strains

Changes in gene expression patterns play a key role in adaptive divergence of strains, helping to shape phenotypic change (Price et al. 2022). To understand changes to the transcriptional landscape in the industrial lineage of *S. clavuligerus,* comparative transcriptomics of the strains was undertaken. RNA was harvested during exponential phase of growth for all strains and their transcriptomes were compared (**Fig. 5A**). A total of 37 genes were found to change expression across all strains (**Supp. Table 10).** To facilitate detailed analysis, all genomes were reannotated with PROKKA (Seemann 2014) and each gene (6124 CDSs) was assigned a corresponding KEGG number where possible (2356/6124, 38.47%).

**Fig. 5.**
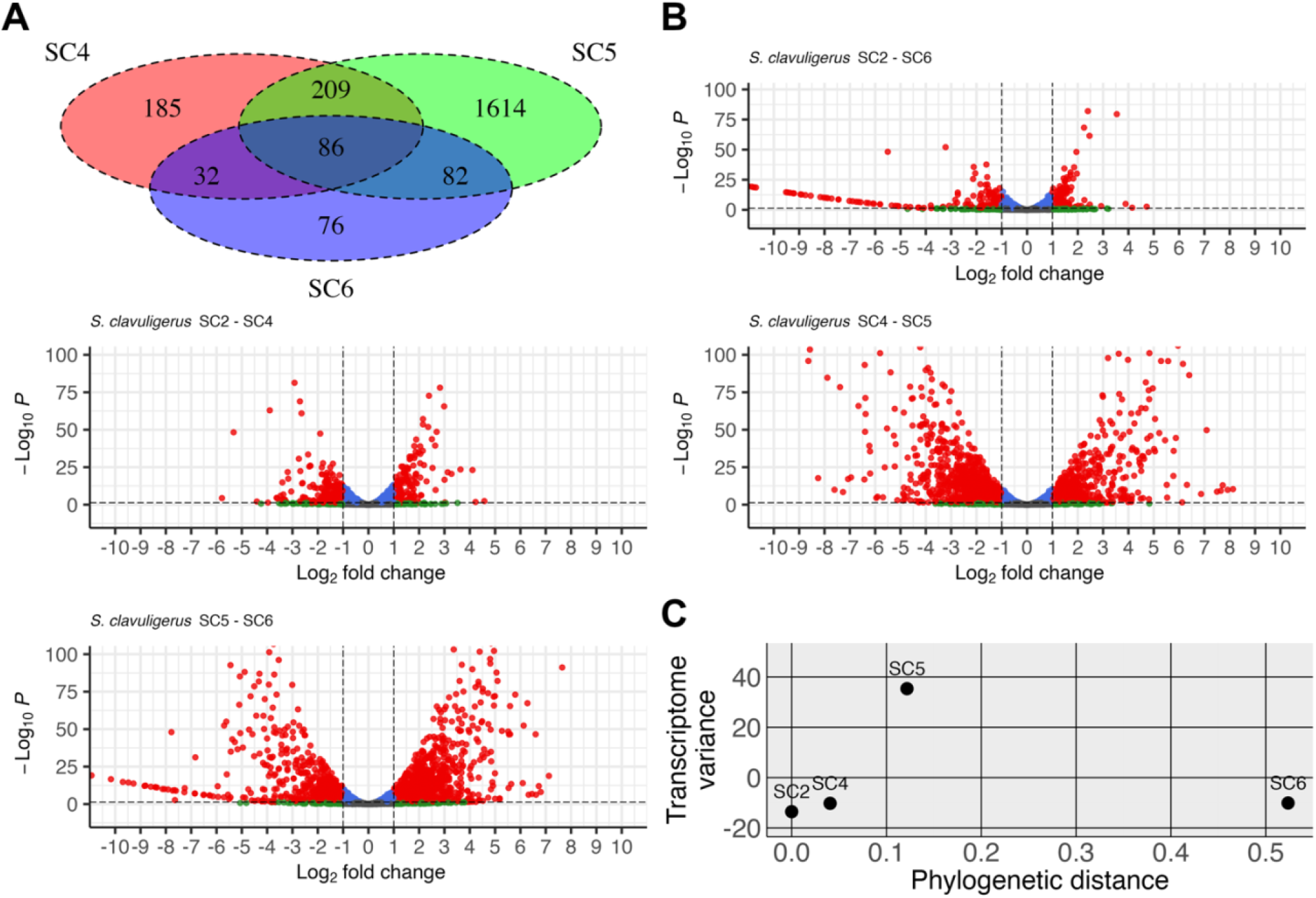
Differential gene expression of *Streptomyces clavuligerus* strains in the industrial lineage. RNA-seq compared for *S. clavuligerus* SC4 to SC6 against the progenitor strain *S. clavuligerus* SC2. **A).** Venn-diagram shows the number of genes with significant changes in expression and how these DEGs map across the lineage. **B).** Volcano plots show the same data with dashed lines indication thresholds for DEGs (DEG significance defined as p-value =<0.05 for 2-fold change, i.e. log2(fold change) threshold +/-1). **C).** Transcriptional variance plot represents the overall variance in the transcriptomes of SC4-SC6 (y-axis) compared to SC2 with the phylogenetic distance (x-axis) of the lineage members. Data points represent the transformed whole transcriptome data reduced to a single data point via a principal component analysis, the data from a single axis (using PC1) was then averaged for the replicate data and plotted against phylogenetic distance.

Transcriptomes of each *S. clavuligerus* strain were compared to the progenitor strain (SC2; **Fig. 1**) to identify differentially expressed genes (DEGs; with the threshold being defined as p-value =<0.05 and log2(fold change) +/- 1); all RNA-seq results can be found in **Supp. Table 4-12**. The DEGS from SC2-SC4 (Supp. Table 4 and 6), SC2-SC5 (Supp. Table 4 and 11) and SC2-SC6 (Supp. Table 4 and 9) are visualised as a venn diagram (**Fig. 5A**). *S. clavuligerus* SC4 had a total of 512 DEGs when compared to *S. clavuligerus* SC2 (**Supp. Table 4 & 6**). *S. clavuligerus* SC5 showed the biggest change in gene expression when compared to its direct progenitor strain (SC4) with 1727 DEGs (**Supp. Table 4 & 7**). Remarkably, *S. clavuligerus* SC6 shows 1770 DEGs when compared to SC5 (**Supp. table 4 & 8**). Most changes observed appear to reverse the changes in expression that arose between SC4-SC5 indicating a functional reversion of the transcriptome to an expression profile much closer to *S. clavuligerus* SC2. This is likely the result of the regulatory gene mutations observed in the strains between *S. clavuligerus* SC5 and SC6. To further investigate this, a comparison of *S. clavuligerus* SC2 to SC5 (**Supp. Table 4 & 9**) and SC2 to SC6 (**Supp. Table 4 & 10**) showed that the majority of DEGs associated with the *S. clavuligerus* SC2 to SC5 comparison appear to be absent in the *S. clavuligerus* SC2 to SC6 comparison. This suggests that the regulatory mutations occurring in the lineage between *S. clavuligerus* SC5 and SC6 may be compensatory in terms of regulating strain physiology, while are also reflected phenotypically in the recovery of catabolic breadth in *S. clavuligerus* SC6 (**Fig. 5B**). To gain an overall picture of how the transcriptome varies between the strains the transcriptional variance was plotted against the phylogenetic distance of the strains SC2-SC6; **Fig. 5C**). The largest transcriptional variance was observed between *S. clavuligerus* SC4-SC5 and SC5-SC6. These observations further support the phenotypic data from Biolog (**Fig. 4A & B)**. A relatively small number of mutations lead to the dramatic difference between SC4-SC5, whereas a larger number of mutations occurred between *S. clavuligerus* SC5-SC6, with the observable recovery of both function (catabolic breadth) and restoration of the transcriptional landscape.

To understand potential impacts of these transcriptional changes in specific metabolic pathways, the expression of genes associated with the biosynthesis of clavulanic acid, cephamycin and the paralogous clavam cluster (located on pSCL4) were investigated. Comparison *S. clavuligerus* SC2 to SC4 shows increased CA production, with some genes in the CA BGC exhibiting reduced gene expression profiles. One exception to this is the increased expression of *oat2,* which encodes an ornithine acetyltransferase (homologue of *argJ -*see below), that is believed to contribute to arginine provision during the early stages of biosynthesis (Fuente et al. 2004; Tahlan et al. 2004). Down regulation of most of the CA BGC was found to occur between the *S. clavuligerus* SC4 and SC5 strains. Whereas these changes return to previous expression levels between *S. clavuligerus* SC5 and SC6 strains to resemble those in *S. clavuligerus* SC2 (**Supp. Fig. 12**). These data suggest that the increase in CA titre in *S. clavuligerus* is unlikely to be due to direct changes in biosynthetic gene expression. Examining the paralogous, but incomplete CA BGC located on pSCL4, indicates a greater difference in gene expression as the lineage of strains advances, with significant up regulation of the early CA biosynthetic genes present in the paralogous BGC between *S. clavuligerus* SC4 and SC5 (**Supp. Fig. 13).** These data suggest that enhanced CA titre may be the result of increased expression of the paralogous partial CA BGC on pSCL4. It is worth noting that the paralogous CA cluster is not in the region of reduced coverage on pSCL4 on *S. clavuligerus* SC5. The cephamycin C BGC is co-located and co-regulated with the CA BGC through the action of CcaR (located in the cephamycin C BGC; (Santamarta et al. 2002). Transcriptional analysis of cephamycin C BGC exhibits a broadly similar pattern of gene expression as CA (**Supp. Fig. 14)** suggesting that simple changes to BGC expression is not responsible for increased industrial CA production.

Despite the similarity in chemical structure (ß-lactam ring), cephamycin C and CA are derived from different starter precursors in *S. clavuligerus* (Liras 1999; Paradkar 2013). To understand the role played by precursor supply in CA biosynthesis the metabolic pathways generating CA precursors was investigated. Pathways generating glyceraldehyde 3-phosphate (G3P; **Supp. Fig. 15**) and arginine were examined (**Supp. Fig. 16**). Genes associated with G3P synthesis were largely unaffected across the lineage, with only the transketolase paralog (SCLAV_1133) exhibiting reduced transcription in SC6, potentially modulating cellular G3P levels between glycolysis and the pentose phosphate pathway (PPP). Arginine biosynthetic genes are all upregulated between *S. clavuligerus* SC2 and SC4, although this appears to have been stabilized in later strains, including *argJ,* the homolog of *oat2* present in the CA BGC and pSCL4-encoded parologous CA BGC.

To further understand how metabolism is modulated during domestication, the major biochemical control points of carbon metabolism were examined (**Supp. Fig. 17**). The major control point for glycolysis is phosphofructokinase, of which *Streptomyces* generally carry three paralogous genes (Borodina et al. 2008; Schniete et al. 2018). These genes show limited changes in expression between the strains. The fructose 1,6 biphosphatase gene (*glpX*), which catalyses the reverse reaction of phosphofructokinase contains a nonsense mutation (W39*; see below) in the lineage from *S. clavuligerus* SC5 onwards, suggesting gluconeogenesis is blocked at this point. The genes encoding enzymes that act as major control points for the tricarboxylic acid (TCA) cycle (Pyruvate dehydrogenase, isocitrate dehydrogenase and a-ketoglutarate decarboxylase), gluconeogenesis (Fructose 1,6 biphophatase II (*glpX*) and pyruvate carboxylase) and pentose phosphate pathway (PPP; Glucose 6 phosphate dehydrogenase) exhibit no change in expression between *S. clavuligerus* SC2 and SC6.

Examining primary metabolism more widely through analysis DEGs (+/- 1; log2 scale and p-value < 0.05), changes were observed in other key central metabolic pathways. *S. clavuligerus* SC2 and SC4 shows 512 DEGs genes with 246/512 hypothetical protein annotations (**Supp. Table 4 and 6**). The KEGG annotated gene sets exhibit 141/217 genes up-regulated and 76/217 down-regulated genes. The genes include 13 encoding proteins associated with ribosomal function, those involved in amino acid biosynthesis (24 DEGs) with many associated with arginine biosynthesis (**Supp. Fig 19 & 20**). There are 18 genes involved in carbon metabolism (**Supp. Fig 21**), including up-regulation of GAPDH (SCLAV_SC2_26880) providing a direct link with a CA precursor molecule, previously targeted by Li & Townsend (2006) for metabolic engineering. Up-regulated carbon metabolic genes in this part of the lineage also shows changes in the malate synthase and isocitrate lyase gene expression suggesting the glyoxylate shunt is active.

In the *S. clavuligerus* lineage step from SC4 to SC5, there are 1728 DEGs with 884/1728 hypothetical protein annotations (**Supp. Table 7**). KEGG annotated genes are 256/677 up-regulated and 421/677 genes down-regulated. There are 50 regulatory proteins included in the DEGs list, along with 39 putative ABC transporters, with four of the up-regulated transporters annotated as being involved in the uptake of glutamate. Remarkably, in *S. clavuligerus* SC5 there is an obvious down-regulation of much of the arginine biosynthesis pathway (**Supp. Fig 22 & 23**), with the pathways that feed in and out of the G3P node of metabolism not exhibiting a consistent trend (**Supp. Fig 24**). There also appears to be a widespread transcriptional shift in expression of purine and pyrimidine biosynthesis, along with genes associated with DNA repair function suggesting general stress responses are active in *S. clavuligerus* SC5 (**Supp.Table 4, 7, 8, & 9**).

DEGs in the late strains of *S. clavuligerus* SC5 to SC6 number 1770 CDSs (**Supp. Table 8**). Examination of these DEGs show that 652/1770 have KEGG annotations, with 944/1770 being annotated as hypothetical proteins. KEGG annotated genes include 376/652 up-regulated genes and 276/652 down-regulated genes. There are 58 putative regulatory proteins that are differentially expressed between the strains, suggesting that regulatory changes are key to driving the enhanced CA production phenotype. Regulatory proteins related to carbon metabolism (**Supp. Fig 25**), multiple steps of glycolysis, the pentose phosphate pathway and in wider pathways leading to the G3P and pyruvate nodes of metabolism are up regulated. There are several changes in ABC transporter expression between *S. clavuligerus* SC5 to SC6 despite the strains being grown on the same medium (**Supp. Fig 26).** Genes encoding transporters for sorbitol/mannitol, glucoside, thiamine, branched chain amino acids, xylose, chitobiose and biotin were up-regulated, whereas those associated with glutamate and galactose were down-regulated. The upregulation of chitiobiose (N-acetyl-glucosamine metabolism) and down regulation of glutamate are supported by the Biolog data (**Fig. 4B).** Changes in amino acid biosynthesis (**Supp. Fig 27**) suggests significant alteration to nitrogen metabolism with glutamine production, the major nitrogen pool within cells, being downregulated. There is up-regulation of fatty acid catabolism which likely reflects the selection bias as oil and substrates such as soya flour are used widely in industrial media for antibiotic production and there are also signals for positive selection in genes associated with beta-oxidation pathways (Petkovic et al. 2006; **Supp. Fig 28 & 29**).

To develop a broader view of the transcriptional changes between the starting strain and the late strain, the transcriptome of *S. clavuligerus* SC2 was compared to SC6. There are 277 DEGs between these two strains (**Supp.Table 9)**, of which 100/277 have KEGG annotations. A total of 41/100 KEGG annotated genes are up-regulated and 59/100 of these genes are down-regulated. Examination of the annotations show that seven of these DEGs are putative regulatory proteins. The NADH dehydrogenase subunits required for oxidative phosphorylation are also downregulated in *S. clavuligerus* SC6 compared to SC2 (**Supp. Fig. 30**). Up-regulation of the glyoxylate shunt genes is clear again and likely reflects the provision of acetyl-CoA for the TCA cycle, potentially an adaptation in these strains selected due widespread use of oil in industrial fermentation media. Most genes in the endiyne natural product, clavulyne (Han et al. 2023) BGC are also downregulated in *S. clavuligerus* SC6 compared to SC2.

### *In trans* expression of genes from the progenitor strain in advanced industrial strains can restore ancestral function

The development of the *S. clavuligerus* lineage has resulted in the accumulation of mutations in successive strains, and phenotypic analysis indicates that CA titre has been altered along with concomitant changes to catabolic capability in the strains. Whilst these mutations on the whole exhibit signals of positive selection, none of the mutations localise to genes directly involved in CA biosynthesis. It was hypothesised that the mutations present in the strains represent adaptive mutations that enhance CA production rather than neutral or deleterious mutations that have few consequences for the strains in the industrial process. To test this hypothesis four genes were selected for complementation *in trans* of the genetic lesions in *S. clavuligerus* SC6 with the progenitor allele of each gene. Each allele representing the *S. clavuligerus* SC2 and SC6 variant of each gene was also introduced into the progenitor strain *S. clavuligerus* SC2. The genes selected comprised different GO groupings, SCLAV_0130, a putative helix-turn-helix containing DNA binding/regulatory protein of unknown function, containing a missense mutation (832C>T; Arg278Trp); SCLAV_1550, a conserved hypothetical protein of unknown function with a missense mutation (550A>G; Ile184Val); SCLAV_1980, a conserved putative amino acid ABC transporter containing missense mutation (155G>A; Arg52His); and a nonsense mutation (117G>A; Trp39*) in the gluconeogenic enzyme, fructose 1,6 biphosphatase gene, *glpX* (**Fig. 6A**).

**Fig.6.**
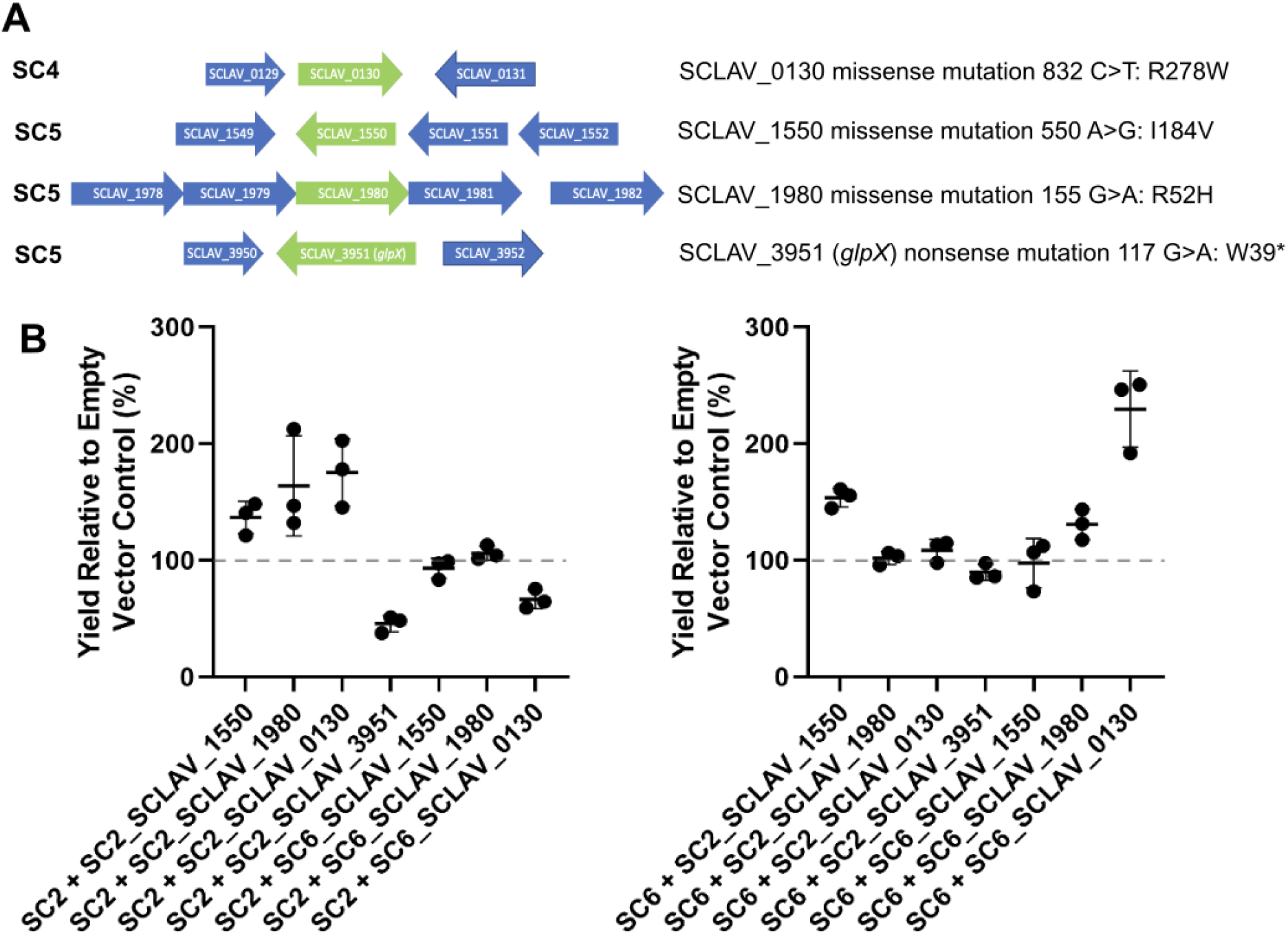
Expression of mutant loci *in trans* in advanced industrial strains of *Streptomyces clavuligerus.* **A)**. Genetic organisation of the genes that were expressed *in trans* in *S. clavuligerus* SC2 and SC6 on the integrating plasmid pIJ10257 and the nature of the mutation. The Strain number SCx refers to the strain in which the mutation arose in the lineage. **B).** Normalised production of CA from *S. clavuligerus* SC2 and SC6 strains containing the additional progenitor and advanced strain mutant alleles grown in TSB. Data represent the mean of three replicates and the error bars represent the standard deviation of the data. Data for the null allele of SCLAV_3951 from *S. clavuligerus* SC6 are not presented as this would represent complementation of the WT allele with a null allele.

Each allele was cloned under the control of the constitutive *ermE\** promoter into the integrating vector pIJ10257 (Hong et al. 2005) and introduced into *S. clavuligerus* SC2 and SC6. The CA production of each strain was compared to the empty vector-containing parental strain in growth on TSB medium. When grown in TSB, the additional copies of the progenitor alleles in *S. clavuligerus* SC2 give rise to an increase in CA production compared to the empty vector control strain likely as the result of gene dosage effects (**Fig. 6B**). The exception to this was an additional copy of *glpX*, which results in a decrease in CA production. It is hypothesised that the additional *glpX* activity results in metabolic flux away from G3P synthesis in this strain. The addition of *S. clavuligerus* SC6 alleles to *S. clavuligerus* SC2 largely results in strains performing less well in terms of their CA production in TSB. When the progenitor strain (*S. clavuligerus* SC2) alleles are introduced to *S. clavuligerus* SC6, only SCLAV_1550 (encoding a hypothetical protein) shows an increase in CA production. Additional copies of the *S. clavuligerus* SC6 alleles in a *S. clavuligerus* SC6 strain background do not increase the production of CA except for the putative DNA-binding protein encoding gene SCLAV_0130. These data suggest that positive and negative epistasis is occurring across the industrial strain lineage in terms of the effects on CA titre, with several mutations in *S. clavuligerus* SC6 unlikely to advance CA titre directly. Moreover, these data suggest that many of the mutations in these strains are neutral in terms of their effect on CA titre.

## Discussion

Domestication of microorganisms was largely accidental until the emergence of genetics as an experimental science in the 1940s. This allowed deliberate introduction of mutations into populations followed by selection of the desired traits such as improving the productivity of industrial cultures (Adrio and Demain 2006). This approach has been very successful in shifting the proportion of favoured products in fermentation broths and increasing production of natural products such as antibiotics (Jeyachandran et al. 2024). Mutational analysis of industrial antibiotic producing strains to date has largely been limited to genomic analysis of high-producing strains compared to a wild-type of the same species. This work has revealed the nature of mutation accumulation in industrial strains within the same lineage, delivering insight into nature and order of the mutations that drive strain evolution over time.

*Streptomyces* are unusual amongst bacteria, possessing a large linear chromosome (6-12 Mb) ended with inverted repeats and an overall high GC content (∼72%; Hoff et al. 2018; Bury-Moné et al. 2023). The genus has generally been considered as genetically unstable with numerous mechanisms explaining the emergence of 0.1-1% spontaneous mutations per generation in culture (Hoff et al. 2018; Tidjani et al. 2019). Genome circularisation, fusion of chromosome ends, and chromosomal arm exchange have been implicated in this genetic instability (Fischer et al. 1998; Uchida et al. 2003; Inoue et al. 2014) and has often hampered industrial strain improvement programs. It is increasingly recognised that plasticity plays a significant role in *Streptomyces* chromosome dynamics and wider biology (Hoff et al. 2018, Gravius et al. 1994; Gomez-Escribano et al. 2021). It is well established that plasmids are an important selective force acting on bacterial genomes, shaping their evolution and core cellular functions (Hall et al. 2024) and there is still much to learn about the role played by the four plasmids in *S. clavuligerus* biology.

The data presented here reinforces the plasticity and dynamics of *Streptomyces* chromosomes, with multiple forms present within the same population and alternative episome integration events being present in the industrial *S. clavuligerus* lineage. These data further highlight the dynamics of bacterial genomes where rearrangements and multiple forms are the norm within populations rather than the exception (Page et al. 2020). No large chromosomal deletions were identified in the *S. clavuligerus* industrial lineage, where in some strains these have been associated with altered antibiotic production (Hoff et al. 2018; Pšeničnik et al. 2024), yet we do see although significant changes to pSCL4.

Random mutagenesis for strain improvement generally leads to rapid improvement in the desired traits (Adrio and Demain 2006), where the mutagenesis programme generates the standing variation within the populations for artificial selection to act, which in this case results in an increase in CA production. When mutation is driving adaptation (or via artificial selection) relatively few mutations are required to deliver large increases in fitness (Wagner 2023). In such ‘adaptive walks’ the advantages gained for a particular trait tend to be successively smaller with each generation (Lenski and Travisano 1994; Saxer et al. 2010; Grant et al. 2021; Favate et al. 2023). This is widely observed in natural selection and has been documented in strain improvement programmes, where the frequency of improved production mutants appearing during selection decreases with each subsequent round of mutagenesis (Adrio and Demain 2006; Jeyachandran et al. 2024).

This study is the first to show accumulation of mutations across an industrial antibiotic producing lineage and assess the phenotypic and genetic consequences of the mutations that become fixed in the production strain. It is also well established in experimental evolution studies that adaptation and concomitant gains in fitness are more rapid in the early stages (Lenski 2017) and this study reinforced this concept in terms of rapid CA production gains in the early stages of the strain improvement process. The nature of selection on industrial media is such that the strains climb a fitness peak under a defined condition, and this can reduce metabolic flexibility of strains, reducing fitness in different conditions (antagonistic pleiotropy). This can be restrictive to industrial processes if modifications to medium or growth conditions are later required to modify media or process changes. This is analogous to adaptation in the natural environment, where fitness gains in a single environment are likely to lead to trade-offs in fitness in alternative environments (Basan et al. 2020; Nguyen et al. 2021; Zhu and Dai 2024). This study provides an example in terms of growth, with the industrial strains becoming more streamlined in their catabolic capability and the production of CA dropping in sub-optimal media. The absence of selective sweeps, as would be the case in the natural environment, serves to fix mutations in the population in stable environment (artificial selection on a single trait) and can result in the fixation of potentially negative mutations within the population. The fixation of the *glpX* nonsense mutation in this *S. clavuligerus* lineage is a clear example of enhanced CA production through metabolic reprogramming through the loss of a metabolic route that redirects flux from CA precursor metabolite.

Regulatory mutations often dominate in experimental evolutional studies (Heyde et al. 2021) and this is also likely in strain improvement programmes, where minimal genetic modifications offer maximum effects in terms of strain improvement (Liu et al. 2013). Detailed experimental studies suggest that overall evolution via regulatory mutations generally does not occur more frequently, but it is likely to have broader effects (Liu et al. 2013; Metzger et al. 2016; Duveau et al. 2021; Stoltzfus 2021).

Complex phenotypic changes are often engineered via random mutagenesis (Bassalo et al. 2018). While other studies in *Streptomyces* strains with enhanced antibiotic production profiles have shown changes within the BGCs, here we have identified mutational changes that act on precursor supply for CA biosynthesis. Condensation of arginine and G3P is the first step of CA biosynthesis (Li and Townsend 2006; Paradkar 2013) and the mutagenesis that generated *S. clavuligerus* SC4 from the progenitor strain (*S. clavuligerus* SC2) exhibits increased expression of genes associated with arginine metabolism. Generally antibiotic production media are carbon rich and regulation of natural product biosynthesis is in response to nitrogen or phosphorus limitation (Fernández-Martínez and Hoskisson 2019). Transcriptional changes broadly indicate an upregulation of arginine metabolism specifically but also more widely amino acid transport suggesting strains are scavenging nitrogen from the environment. Modifications to carbon metabolic genes mostly are associated with anaplerosis and gluconeogenesis (glyoxylate shunt genes and *glpX*) suggesting that much of the carbon demand for CA through G3P is not supplied via glycolysis. Previous work was able to show that deletion of one glyceraldehyde-3-phosphate dehydrogenase paralogue enhanced CA production only when arginine was supplied exogenously (Li and Townsend 2006).

Increased knowledge of *Streptomyces* strain improvement, domestication and metabolic modifications may inform of the development of future industrial strains, and how strain improvement processes and mutational landscapes may constrain our future ability to modify industrial processes to modify feedstock with a view to using more sustainable carbon sources.

## Materials and Methods

### Growth and Maintenance of Bacterial Strains

The *Streptomyces clavuligerus* strains used in this study are detailed in **Table 1**. The genetic constructs used are summarised in **Table 1**. Routine growth and maintenance procedures were carried out according to (Feeney et al. 2022) for 10 days for preparation of spores. Liquid cultures were routinely grown in 250 mL Erlenmeyer flasks in Tryptone soy broth (TSB; Oxoid), using a two-day pre-culture to ensure consistency of inoculum between replicate flasks. Briefly, *S. clavuligerus* spore stocks (1 x 10^8^ spores) were used to inoculate a 50 mL seed culture of TSB which was incubated at 26 °C with shaking at 250 rpm. After 48 hrs, cells were centrifuged at 4000 x *g* and resuspended at an OD_600nm_ of 0.4 and 2 mL inoculated into fresh medium (∼2 x 10^6^ CFUs).

**Table 1.**
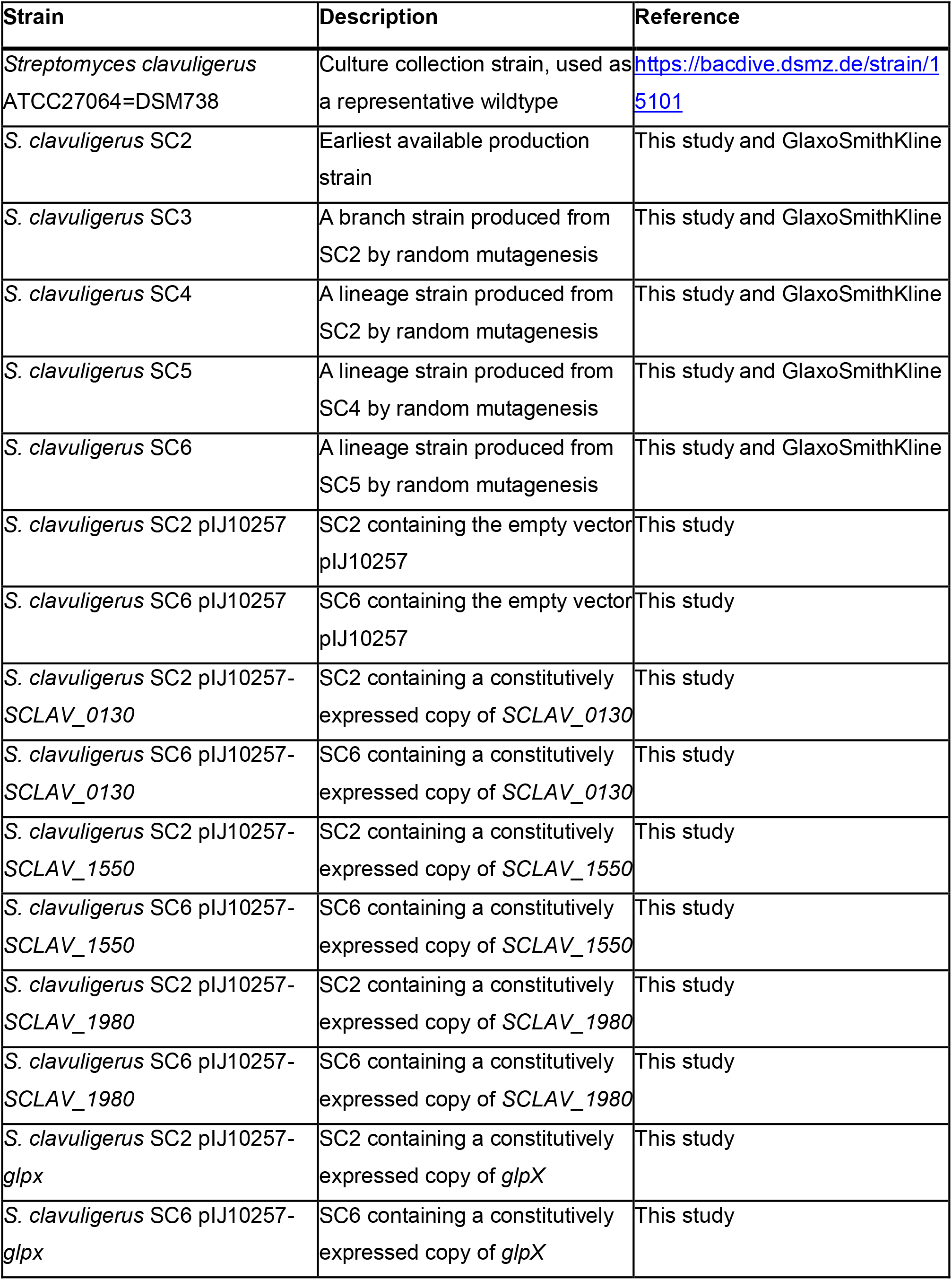
Strains Used in this study.

**Table 2.** Plasmids used in this study.

| Plasmid | Description | Reference |
| --- | --- | --- |
| pIJ10257 | *ermEp\** with ribosome binding site cloned into pMS81. Hygromycin resistant/ϕiBT1 intergrase and *att* site. | Hong et al., 2005 |
| pIJ10257-SCLAV_0130 | pIJ10257 with *SCLAV_0130* cloned under the control of *ermEp\** | This study |
| pIJ10257-SCLAV_1550 | pIJ10257 with *SCLAV_1550* cloned under the control of *ermEp\** | This study |
| pIJ10257-SCLAV_1980 | pIJ10257 with *SCLAV_1980* cloned under the control of *ermEp\** | This study |
| pIJ10257-*glpX* | pIJ10257 with *glpX* cloned under the control of *ermEp\** | This study |

### DNA extraction and Genome sequencing

Genomic DNA was extracted from *S. clavuligerus* using the *Streptomyces* DNA isolation protocol described by (Feeney et al. 2022)*. S. clavuligerus* SC2 through to SC6 were grown in 250 mL Erlenmeyer flasks in 50 mL Tryptone soy broth (TSB; Oxoid) from ∼1x 10^8^ spores at 26 °C with shaking at 250 rpm. After 48 hrs, cultures were centrifuged at 4000 x *g,* supernatant removed and cells resuspended in 300 µL TE buffer (Tris-HCl 50 mM, EDTA 20 mM) with 100 mg\mL lysozyme and 0.1 mg/mL RNase (Qiagen) and incubated at 30°C for 30 minutes. To each sample, 50 µL 10% SDS was added and mixed, followed by 85 µL of 5M NaCl and mixed. Then, add 10 mL of 20 mg/mL proteinase K and thoroughly mixed. Samples were incubated at 55°C for 2.5 hours. Phenol: chloroform was added (1 volume) to each, vortexed for 15 seconds, followed by centrifugation for 10 minutes at 13,000 rpm. The aqueous phase was removed and combined with 300 µL chloroform, vortexed and centrifuged (10 minutes at 13,000 rpm). The upper aqueous phase was removed and precipitated by adding 500 µL of 100% isopropanol. Samples were centrifuged, the supernatant removed and washed with 1 mL of cold, 70% ethanol. Supernatant was removed, pellet briefly dried and resuspended in DNase free water. Short-read data was obtained from two sources: lllumina MiniSeq instrument using Nextera DNA Flex Library Preparation kit with samples pooled evenly for SC2-SC5 and, for SC6 data, using an Illumina NovaSeq 6000 by Novogene. PacBio sequencing and assembly of *S. clavuligerus* SC6 was provided by The University of Liverpool and the Earlham institute, with the same procedures described in (Gomez-Escribano et al. 2021). PacBio sequencing of *S. clavuligerus* SC2 was provided by Nu-omics (University of Northumbria, UK) using the PacBio Sequel instrument. In brief, high quality gDNA (>1.8 A260/A280 ratio) was size sheared using a Covaris g-Tube to ∼9 kb assessed by Agilent Bioanalyzer gel. Samples were treated with ExoVII to remove ssDNA and fragment ends were repaired to make blunt end dsDNA. Samples were ligated to SMRTbell adapter hairpins and treated with ExoIII to remove failed ligations. For each of these steps, AMPure magnetic beads were used for clean-up. Sequencing primers (v3) were annealed to the SMRTbell templates. PacBio Polymerase binding reaction to SMRTbell template was carried out via the Sequel Binding kit, V2.1 before sequencing on the PacBIo Sequel instrument. Samples were sequenced using 10 h movie capture.

A combination of approaches to data processing were used. All PacBio data were assembled using HGAP4 (Chin et al. 2013). All Illumina data were assembled *de novo* using SPAdes and SC2 and SC6 mapped to their respective PacBio assemblies (Scripts available at https://github.com/PaulHoskisson/industrial-lineage-of-S.clavuligerus). Additionally, hybrid assemblies using a combination of Illumina and PacBio data were carried out using SPAdes (Bankevich et al. 2012). These were then used as the base for manually stitching the genomes together and, where relevant identification of multiple forms was required, these were confirmed by PCR and sequencing. All sequence data are deposited at Sequence Read Archive (SRA) SAMN42008322-6. Reannotation of the *S. clavuligerus* genome was created using Prokka (Seemann 2014) to ensure consistency across the lineage and can be accessed at the GenBank Bioproject PRJNA1127551.

Genome annotation was performed using Prokka version 1.14.6 (Seemann 2014) and the amnio acid sequences used to generate KEGG identifiers using BlastKOALA version 3 (Kanehisa et al. 2016).

### Comparative genomics and SNP calling

Single Nucleotide Change calling was carried out using SNIPPY (https://github.com/tseemann/snippy; Version 4.6.0) using default settings and the SC2-1 genbank file as the reference sequence (https://github.com/PaulHoskisson/industrial-lineage-of-S.clavuligerus/blob/main/01_Comparitive_genomics_and%20SNP_calling/SC2-1.gbk). The resulting CSV files were combined into a single file using an R script (https://github.com/PaulHoskisson/industrial-lineage-of-S.clavuligerus/blob/main/01_Comparitive_genomics_and%20SNP_calling/03_snippy_combine.R). The resulting table can be found in **Supp. Table 2**.

### Structural comparison of *Streptomyces clavuligerus* genomes

The nucleotide synteny plots for all Prokka annotated genomes were generated using the PyGenomeViz Python package (https://moshi4.github.io/pyGenomeViz/), with genomes aligned with NUCmer v3.1 ((Kurtz et al. 2004) using many-to-many alignments.

### Omnilog Biolog analysis

Catabolic capability of the *S. clavuligerus* strains was determined using OmniLog analysis with Biology plates PM1 (Biolog, 12111). Strains were grown from spore stocks on plates for two days at 26°C. Three representative single colonies were selected and streaked onto fresh L3M9 for lawns. After a further two days growth, biomass was scraped from the plates and added to inoculation fluid IF-0a (Biolog 72268) to an O.D_600_ of 0.04, to create the initial cell suspension. Experimental inoculation fluid (per 24 mL) was prepared using 20 mL stock bottle solution of IF-0a, plus addition of 1.2 mL metal ion cocktail (containing 5 mM each: ZnCl_2_ 7H_2_O, FeCl_2_ 6H_2_O MnCl_2_ 4H_2_O, CaCl_2_ 2H_2_O, filter sterilised) 0.24 mL of Dye Mix D (Biolog, 74224), 1.2 mL of sterile dH_2_O and 2.32 mL of cell suspension to a final volume of 24 mL. This was then inoculated into PM1 plates and incubated at 26°C for 7 days. Data were extracted using Biolog software (conversion of D5E to OKA: D5E_OKA Data File Converter v1.1.1.15 and extraction of raw kinetic data using PM analysis software: Kinetic V1.3) and visualised and compared using R scripts (https://github.com/PaulHoskisson/industrial-lineage-of-S.clavuligerus). Data were compared using a one-way ANOVA with a P-value threshold of 0.05. Growth was scored when absorbance was 1.5x growth of negative control (∼50 growth units). Samples were processed using available scripts (https://github.com/PaulHoskisson/industrial-lineage-of-S.clavuligerus/tree/main/02_Omnilog_biolog_analysis). Two approaches were used. The first was to select 48h data and generate growth curves to visually inspect the data. The second was to apply the data to BactExtract(Dénéréaz & Veening 2024) selecting all output and filtering this to select samples with growth.

### RNASeq and analysis

*S. clavuligerus* SC2-SC6 were grown as above. After 48 hrs, cells were centrifuged at 4000 x *g and* resuspended at an OD_600_ of 0.4 and 2 mL inoculated into fresh medium. Samples were grown to exponential phase (∼48 hrs). Samples (1 mL of culture) were centrifuged at max speed (16,160 x g) for 1 minute in 2 ml screw capped tubes and resuspended in 1 mL of RNA-later (∼5 volumes to 100-200 µL of wet biomass) and stored at -20°C until extraction (a minimum of 30 minutes). Samples were centrifuged, supernatant removed and resuspended in 300 µL TE buffer (Tris-HCl 50 mM, EDTA 20 mM) with 100 mg\mL lysozyme and incubated at 30°C for 30 minutes. To each sample, 50 µL 10% SDS was added and mixed, followed by 85 µL of 5M NaCl and mixed. Trizol was added (1 mL) to each, vortexed for 15 seconds and centrifuged for 10 minutes. The aqueous phase was removed and combined with 300 µL chloroform, vortexed and centrifuged. The upper aqueous phase was removed and passed through Qiagen RNeasy kit following the manufacturer’s instructions, including on-column DNase treatment (using Qiagen RNase free DNase). Samples were resuspended in 90 µL RNase free water and an off-column DNase Treatment carried out (following manufacturer’s instructions). Samples were passed through a second round of RNeasy kit extraction minus DNase steps. All samples passed their standard QC and were sequenced by Novogene (Novogene UK Ltd.). Resulting Fastq files were processed, using Salmon (version 1.10.3) and R scripts available at https://github.com/PaulHoskisson/industrial-lineage-of-S.clavuligerus/tree/main/03_RNA_seq. Venn diagrams were produced using draw.quad.venn (https://rdrr.io/cran/VennDiagram/man/draw.quad.venn.html). Heat maps were generated using pheatmap (https://www.rdocumentation.org/packages/pheatmap/versions/1.0.12/topics/pheatmap). KEGG maps where generated using KEGG mapper color (https://www.genome.jp/kegg/mapper/color.html - used July 2024, last update Nov 2023).

There is a master supplementary table containing all unfiltered RNA-seq results with KEGG annotations (**Suppl. Table 4**). For each logical comparison of the RNA-seq results, there is an associated supplementary table (All comparisons, SC2vsSC3, SC2vsSC4, SC4vsSC5, SC5vsSC6 and SC2vsSC6, **Supp. Tables 4-9**), containing results that passed the fold change and p-value thresholds.

### Phylogenetic distance and transcriptome variance

Using the identified SNCs from the chromosome of each genome in the lineage, phylogenetic distances were calculated using Clustalomega (version 1.2.4) using default settings on a string of DNA consisting only of the SNCs (**Supp. File 1**). Transcriptome variance was determined by performing a principal component analysis (PCA - using the DESeq2 function plotPCA) and extracting coordinate data from the first principle (PC1). This was performed in lines 195-286 of our DESeq2 R script (https://github.com/PaulHoskisson/industrial-lineage-of-S.clavuligerus/tree/main/03_RNA_seq/01_Deseq_17_03_25.R). Briefly, counts and colData were extracted from each differential expression analysis and combined into a matrix, with data log transformed before plotting. Coordinates were extracted and the average of each PC1 group (i.e. for each strain) and used to indicate transcriptome variance. Supp. File 1 contains the ClustalOmega input sequences, the output (phylogenetic distance), the PCA analysis plot and the corrdinates of the data used for transcriptome distance (PC1).

### Construction of complementation vectors for reconstitution of carbon metabolism

Plasmid constructs for the *glpX*, SCLAV_0130, SCLAV_1550 and SCLAV_1980 alleles from *S. clavuligerus* SC2 and SC6 were synthesised by Genscript in pIJ10257 (Hong et al. 2005) at HindIII/NdeI sites, allowing for expression under the constitutive *ermE\** promoter.

### Clavulanic acid assay

CA concentrations in culture supernatant were determined according to a method modified from Bird et al., (Bird et al. 1982). A derivatization solution was made from 10 g imidazole (VWR chemicals), 7.2 mL 10M HCl (Fisher Scientific), made up to 100 ml with dH_2_O. In a 96 well plate, 200 µL of the derivatization solution was mixed with 8 µL of potassium clavulanate standards in TSB (Merck) or 8 µL of culture supernatant, clarified by centrifugation at 4,000 g for 10 minutes. Plates were incubated for 30 minutes at room temperature before absorbance was determined at 324 nm using a FlexStation 3 plate reader (VWR). The production of clavulanic acid was determined by dividing CA titre by the dry weight.

## Data repositories

**GenBank Bioproject:** PRJNA1127551, Sequence Read Archive (SRA): SAMN42008322-6 and SAMN51091877

**Gene Expression Omnibus (GEO):** GSE212322

**Supplementary Data Files** - https://github.com/PaulHoskisson/industrial-lineage-of-S.clavuligerus/tree/main/supplmentary_material

Code used - https://github.com/PaulHoskisson/industrial-lineage-of-S.clavuligerus

## Code and availability

All code is available: https://github.com/PaulHoskisson/industrial-lineage-of-S.clavuligerus/.

## Sequence data deposits

All genomic sequence data has been deposited with SRA (SAMN42008322-6) and Genbank with NCBI generated annotations (1127551) – also available with PROKKA generated annotations used in this analysis:

https://github.com/PaulHoskisson/industrial-lineage-of-S.clavuligerus/tree/main/01_Comparitive_genomics_and%20SNP_calling. All transcriptional data has been deposited with GEO (GSE212322).

## CRediT (Contributor roles taxonomy)

Conceptualization – PAH.

Data curation - JTM, DEL, REM, JB, KR, JTC, ABK, JPG-E

Formal analysis - JTM, DEL, REM, JB, KR, JTC, ABK, PAH, JPG-E.

Funding acquisition – BW, PAH.

Investigation - JTM, DEL, REM, JB, KR, JPG-E

Methodology - JTM, ISH, BDH, SGK, PAH, BW, JPG-E.

Project administration - JTM, PAH, BW

Supervision - JTM, ISH, PAH, BW.

Validation - JTM, DEL, REM, JB, KR, JTC, ABK, JPG-E, PAH.

Visualization – JTM, DEL, REM, JB, KR, JTC, ABK, PAH.

Writing – original draft – PAH.

Writing – review & editing - JTM, DEL, REM, JB, KR, JTC, ABK, JPG-E, NAC, AJC, SGK, BDH, BW, ISH, PAH.

All authors gave final approval for publication and agreed to be held accountable for the work performed therein.

## Acknowledgements

We would like to thank Prof. Daniel Wilson for assistance in running Genomegamap.

## Supplementary Data Files

**Supplementary Table 1.** Oligonucleotide sequences used during genome structural analysis.

**Supplementary Table 2.** Combined SNCs from SC2-SC6.

**Supplementary Table 3.** Gene ontology results.

**Supplementary Table 4.** Master table of unfiltered RNA-seq DEGs.

**Supplementary Table 5.** List of SC2-SC3 DEGs (130), subset of Supp. Table 4.

**Supplementary Table 6.** List of SC2-SC4 DEGs (370), subset of Supp. Table 4.

**Supplementary Table 7.** List of SC4-SC5 DEGs (1390), subset of Supp. Table 4.

**Supplementary Table 8.** List of SC5-SC6 DEGs (1449), subset of Supp. Table 4.

**Supplementary Table 9.** List of SC2-SC6 DEGs (158), subset of Supp. Table 4.

**Supplementary Table 10.** List of SC2-All DEGs (37), subset of Supp. Table 4.

**Supplementary Table 11.** List of SC2-SC5 DEGS (1991), subset of Supp. Table 4.

**Supplementary Table 12.** List of SC2-All DEGS, excluding SC3 (86), subset of Supp. Table 4.

**Supplementary Figure 1.** PCR reactions of SC2-SC5, Chromosome gap 1.

**Supplementary Figure 2.** PCR reactions of SC2, pSCL3 and chromosome rearrangement.

**Supplementary Figure 3.** SC2-SC5, PCR reactions of confirmation of intact chromosome over region of pSCL3 rearrangement.

**Supplementary Figure 4.** PCR reactions of SC2-SC5, confirmation of intact pSCL1 (Right end).

**Supplementary Figure 5.** PCR reactions of SC2-SC5, confirmation of intact pSCL1 (middle).

**Supplementary Figure 6.** PCR reaction of SC2-SC5, pSCL3 gap.

**Supplementary Figure 7.** PCR reactions of SC6 confirmation of intact pSCL1.

**Supplementary Figure 8.** PCR reaction of Confirmation of SC6 pSCL1 merge to pSCL4.

**Supplementary Figure 9.** PCR reactions of Confirmation of SC6 pSCL4 merge to chromosome merge.

**Supplementary Figure 10.** PCR reactions of Confirmation of SC6 intact chromosome (at site of pSCL4 merge).

**Supplementary Figure 11.** Graphical representation of the SC6.1 and SC6.2 genomic rearrangements.

**Supplementary Figure 12.** RNA-seq results (log2 fold change) for the clavulanic acid biosynthetic gene cluster represented as a heatmap comparing lineage strain expression levels.

**Supplementary Figure 13.** RNA-seq results (log2 fold change) for the cephamycin biosynthetic gene cluster represented as a heatmap comparing lineage strain expression levels.

**Supplementary Figure 14.** RNA-seq results (log2 fold change) for the paralogous and partial clavulanic acid biosynthetic gene cluster represented as a heatmap comparing lineage strain expression levels.

**Supplementary Figure 15**. RNA-seq results (log2 fold change) for the d-glyceraldehyde 3-phosphate (G3P) pathway represented as a heatmap comparing lineage strain expression levels.

**Supplementary Figure 16.** RNA-seq results (log2 fold change) for the L-arginine biosynthetic pathway represented as a heatmap comparing lineage strain expression levels.

**Supplementary Figure 17.** RNA-seq results (log2 fold change) for major control points in carbon metabolism as a heatmap comparing lineage strain expression levels.

**Supplementary Figure 18.** Altered carbon metabolism between SC2 and SC3 using significant DEGs with KEGG annotations (5).

**Supplementary Figure 19.** An overview of SC2-SC4 DEGs from RNA-seq that impact amino acid biosynthesis.

**Supplementary Figure 20.** An overview of SC2-SC4 DEGs from RNA-seq that impact arginine biosynthesis.

**Supplementary Figure 21.** Altered carbon metabolism between SC2 and SC4 using significant DEGs with KEGG annotations.

**Supplementary Figure 22.** An overview of SC4-SC5 DEGs from RNA-seq that impact amino acid biosynthesis.

**Supplementary Figure 23.** An overview of SC2-SC4 DEGs from RNA-seq that impact arginine biosynthesis.

**Supplementary Figure 24.** Altered carbon metabolism between SC4 and SC5 using significant DEGs with KEGG annotations.

**Supplementary Figure 25.** Altered carbon metabolism between SC5 and SC6 using significant DEGs with KEGG annotations.

**Supplementary Figure 26.** Altered ABC transporters between SC5 and SC6 using significant DEGs with KEGG annotations.

**Supplementary Figure 27.** Altered amino acid biosynthesis between SC5 and SC6 using significant DEGs with KEGG annotations.

**Supplementary Figure 28.** Altered fatty acid metabolism between SC5 and SC6 using significant DEGs with KEGG annotations.

**Supplementary Figure 29.** Altered fatty acid degradation between SC5 and SC6 using significant DEGs with KEGG annotations.

**Supplementary Figure 30.** Altered oxidative phosphorylation between SC5 and SC6 using significant DEGs with KEGG annotations.

**Supplementary File 1.** Additional methods material used for phylogenetic analysis (input sequences), PCA plot and coordinates generated in R-script used for figure 5C.

## References

Adrio JL, Demain AL. 2006. Genetic improvement of processes yielding microbial products. FEMS Microbiol. Rev. 30:187–214.

Baltz RH. 2011. Strain improvement in actinomycetes in the postgenomic era. Journal of Industrial Microbiology & Biotechnology 43:343–370.

Bankevich A, Nurk S, Antipov D, Gurevich AA, Dvorkin M, Kulikov AS, Lesin VM, Nikolenko SI, Pham S, Prjibelski AD, et al. 2012. SPAdes: A New Genome Assembly Algorithm and Its Applications to Single-Cell Sequencing. J Comput Biol 19:455–477.

Basan M, Honda T, Christodoulou D, Hörl M, Chang Y-F, Leoncini E, Mukherjee A, Okano H, Taylor BR, Silverman JM, et al. 2020. A universal trade-off between growth and lag in fluctuating environments. Nature 584:470–474.

Bassalo MC, Garst AD, Choudhury A, Grau WC, Oh EJ, Spindler E, Lipscomb T, Gill RT. 2018. Deep scanning lysine metabolism in *Escherichia coli*. Mol. Syst. Biol. 14:e8371.

Bird AE, Bellis JM, Gasson BC. 1982. Spectrophotometric assay of clavulanic acid by reaction with imidazole. Analyst 107:1241–1245.

Borodina I, Siebring J, Zhang J, Smith CP, Keulen G van, Dijkhuizen L, Nielsen J. 2008. Antibiotic Overproduction in *Streptomyces coelicolor* A3(2) Mediated by Phosphofructokinase Deletion. Journal of Biological Chemistry 283:25186–25199.

Bury-Moné S, Thibessard A, Lioy VS, Leblond P. 2023. Dynamics of the *Streptomyces* chromosome: chance and necessity. Trends Genet. 39:873–887.

Cao G et al. 2016. Complete Genome Sequence of *Streptomyces clavuligerus* F613-1, an Industrial Producer of Clavulanic Acid. Genome Announc. 4:e01020–16

Chevrette MG, Gutiérrez-García K, Selem-Mojica N, Aguilar-Martínez C, Yañez-Olvera A, Ramos-Aboites HE, Hoskisson PA, Barona-Gómez F. 2019. Evolutionary dynamics of natural product biosynthesis in bacteria. Nat Prod Rep 37:566–599.

Chin C-S, Alexander DH, Marks P, Klammer AA, Drake J, Heiner C, Clum A, Copeland A, Huddleston J, Eichler EE, et al. 2013. Nonhybrid, finished microbial genome assemblies from long-read SMRT sequencing data. Nat. Methods 10:563–569.

Cho HS, Jo JC, Shin C-H, Lee N, Choi J-S, Cho B-K, Roe J-H, Kim C-W, Kwon HJ, Yoon YJ. 2019. Improved production of clavulanic acid by reverse engineering and overexpression of the regulatory genes in an industrial *Streptomyces clavuligerus* strain. J Ind Microbiol Biot 46:1205– 1215.

Cisneros-Mayoral S, Graña-Miraglia L, Pérez-Morales D, Peña-Miller R, Fuentes-Hernández A. 2022. Evolutionary History and Strength of Selection Determine the Rate of Antibiotic Resistance Adaptation. Mol. Biol. Evol. 39:msac185.

Darwin C. 1859. The Origin of species. John Murray

Dénéréaz J, Veening J-W. 2024. BactEXTRACT: an R Shiny app to quickly extract, plot and analyse bacterial growth and gene expression data. Access Microbiol. 6:000742.v3

Duveau F, Zande PV, Metzger BP, Diaz CJ, Walker EA, Tryban S, Siddiq MA, Yang B, Wittkopp PJ. 2021. Mutational sources of trans-regulatory variation affecting gene expression in *Saccharomyces cerevisiae*. eLife 10:e67806.

Favate JS, Skalenko KS, Chiles E, Su X, Yadavalli SS, Shah P. 2023. Linking genotypic and phenotypic changes in the *E. coli* long-term evolution experiment using metabolomics. eLife 12:RP87039.

Feeney MA, Newitt JT, Addington E, Algora-Gallardo L, Allan C, Balis L, Birke AS, Castaño-Espriu L, Charkoudian LK, Devine R, et al. 2022. ActinoBase: tools and protocols for researchers working on *Streptomyces* and other filamentous actinobacteria. Microb Genom 8.

Fernández-Martínez LT, Hoskisson PA. 2019. Expanding, integrating, sensing and responding: the role of primary metabolism in specialised metabolite production. Curr Opin Microbiol 51:16–21.

Fiedurek J, Trytek M, Szczodrak J. 2017. Strain improvement of industrially important microorganisms based on resistance to toxic metabolites and abiotic stress. J Basic Microb 57:445–459.

Fischer G, Wenner T, Decaris B, Leblond P. 1998. Chromosomal arm replacement generates a high level of intraspecific polymorphism in the terminal inverted repeats of the linear chromosomal DNA of *Streptomyces ambofaciens*. Proc. Natl. Acad. Sci. 95:14296–14301.

Fuente A de la, Martín JF, Rodríguez-García A, Liras P. 2004. Two Proteins with Ornithine Acetyltransferase Activity Show Different Functions in *Streptomyces clavuligerus*: Oat2 Modulates Clavulanic Acid Biosynthesis in Response to Arginine. J. Bacteriol. 186:6501–6507.

Gomez-Escribano JP, Gallardo LA, Bozhüyük KAJ, Kendrew SG, Huckle BD, Crowhurst NA, Bibb MJ, Collis AJ, Micklefield J, Herron PR, et al. 2021. Genome editing reveals that pSCL4 is required for chromosome linearity in *Streptomyces clavuligerus*. Microb. Genom. 7:000669.

Grant NA, Magid AA, Franklin J, Dufour Y, Lenski RE. 2021. Changes in Cell Size and Shape during 50,000 Generations of Experimental Evolution with *Escherichia coli*. J Bacteriol 203.

Gravius B, Bezmalinović T, Hranueli D, Cullum J. 1993. Genetic instability and strain degeneration in *Streptomyces rimosus*. Appl Environ Microb 59:2220–2228.

Gravius B, Glocker D, Pigac J, Pandza K, Hranueli D, Cullum J. 1994. The 387 kb linear plasmid pPZG101 of *Streptomyces rimosus* and its interactions with the chromosome. Microbiology 140:22712277.

Gregory TR. 2009. Artificial Selection and Domestication: Modern Lessons from Darwin’s Enduring Analogy. Evol Educ Outreach 2:5–27.

Hall RJ, Snaith AE, Thomas MJN, Brockhurst MA, McNally A. 2024. Multidrug resistance plasmids commonly reprogram the expression of metabolic genes in *Escherichia coli*. mSystems:e0119323.

Han EJ, Lee SR, Townsend CA, Seyedsayamdost MR. 2023. Targeted Discovery of Cryptic Enediyne Natural Products via FRET-Coupled High-Throughput Elicitor Screening. ACS Chem. Biol. 18:1854–1862.

Heyde SAH, Frendorf PO, Lauritsen I, Nørholm MHH. 2021. Restoring Global Gene Regulation through Experimental Evolution Uncovers a NAP (Nucleoid-Associated Protein)-Like Behavior of Crp/Cap. mBio 12:e02028–21.

Higgens CE, Kastner RE. 1971. *Streptomyces clavuligerus* sp. nov., a β-Lactam Antibiotic Producer. Int J Syst Evol Micro 21:326–331.

Hoff G, Bertrand C, Piotrowski E, Thibessard A, Leblond P. 2018. Genome plasticity is governed by double strand break DNA repair in *Streptomyces*. Sci. Rep. 8:5272.

Hong H-J, Hutchings MI, Hill LM, Buttner MJ. 2005. The Role of the Novel Fem Protein VanK in Vancomycin Resistance in *Streptomyces coelicolor*. J. Biol. Chem. 280:13055–13061.

Hoskisson PA, Seipke RF. 2020. Cryptic or Silent? The Known Unknowns, Unknown Knowns, and Unknown Unknowns of Secondary Metabolism. mBio 11:e02642–20.

Hwang S et al. 2019. Primary transcriptome and translatome analysis determines transcriptional and translational regulatory elements encoded in the *Streptomyces clavuligerus* genome. Nucleic Acids Res. doi: 10.1093/nar/gkz471.

Inoue S, Higashiyama K, Uchida T, Hiratsu K, Kinashi H. 2014. Chromosomal Circularization in *Streptomyces griseus* by Nonhomologous Recombination of Deletion Ends. *Biosci.*, Biotechnol. Biochem. 67:1101–1108.

Jeyachandran S, Vibhute P, Kumar D, Ragavendran C. 2024. Random mutagenesis as a tool for industrial strain improvement for enhanced production of antibiotics: a review. Mol. Biol. Rep. 51:19.

Kanehisa M, Sato Y, Morishima K. 2016. BlastKOALA and GhostKOALA: KEGG Tools for Functional Characterization of Genome and Metagenome Sequences. J. Mol. Biol. 428:726–731.

Kurtz S et al. 2004. Versatile and open software for comparing large genomes. Genome Biol. 5:R12.

Lenski RE. 2017. Experimental evolution and the dynamics of adaptation and genome evolution in microbial populations. ISME J. 11:2181–2194.

Lenski RE, Travisano M. 1994. Dynamics of adaptation and diversification: a 10,000-generation experiment with bacterial populations. Proc. Natl. Acad. Sci. 91:6808–6814.

Li R, Townsend CA. 2006. Rational strain improvement for enhanced clavulanic acid production by genetic engineering of the glycolytic pathway in *Streptomyces clavuligerus*. Metabolic Engineering 8:240252.

Liras P. 1999. Biosynthesis and molecular genetics of cephamycins. Antonie Van Leeuwenhoek 75:109–124.

Liras P, Martín JF. 2021. *Streptomyces clavuligerus*: The Omics Era. J. Ind. Microbiol. Biotechnol. 48:kuab072.

Liu G, Zhang L, Qin Y, Zou G, Li Z, Yan X, Wei X, Chen M, Chen L, Zheng K, et al. 2013. Long-term strain improvements accumulate mutations in regulatory elements responsible for hyper-production of cellulolytic enzymes. Sci. Rep. 3:1569.

Medema Marnix H, Alam MT, Breitling R, Takano E. 2011a. The future of industrial antibiotic production: From random mutagenesis to synthetic biology. Bioengineered Bugs 2.

Medema Marnix H., Alam MT, Heijne WHM, Berg MA van den, Müller U, Trefzer A, Bovenberg RAL, Breitling R, Takano E. 2011b. Genome-wide gene expression changes in an industrial clavulanic acid overproduction strain of *Streptomyces clavuligerus*. Microb Biotechnol 4:300–305.

Medema MH, Trefzer A, Kovalchuk A, Berg M van den, Müller U, Heijne W, Wu L, Alam MT, Ronning CM, Nierman WC, et al. 2010. The Sequence of a 1.8-Mb Bacterial Linear Plasmid Reveals a Rich Evolutionary Reservoir of Secondary Metabolic Pathways. Genome Biology and Evolution 2:212–224.

Metzger BPH, Duveau F, Yuan DC, Tryban S, Yang B, Wittkopp PJ. 2016. Contrasting Frequencies and Effects of cis- and trans-Regulatory Mutations Affecting Gene Expression. Mol. Biol. Evol. 33:1131–1146.

Nguyen J, Fernandez V, Pontrelli S, Sauer U, Ackermann M, Stocker R. 2021. A distinct growth physiology enhances bacterial growth under rapid nutrient fluctuations. Nat. Commun. 12:3662.

Nielsen J. 1997. Physiological engineering aspects of *Penicillium chrysogenum*. Singapore: World Scientific

Page AJ, Ainsworth EV, Langridge GC. 2020. socru: typing of genome-level order and orientation around ribosomal operons in bacteria. Microb Genom. 6: 10.1099/mgen.0.000396.

Paradkar A. 2013. Clavulanic acid production by *Streptomyces clavuligerus*: biogenesis, regulation and strain improvement. The Journal of Antibiotics 66:411–420.

Petkovic H, Cullum J, Hranueli D, Hunter IS, Peric-Concha N, Pigac J, Thamchaipenet A, Vujaklija D, Long PF. 2006. Genetics of *Streptomyces rimosus*, the Oxytetracycline Producer. Microbiology and Molecular Biology Reviews 70:704728.

Price PD, Droguett DHP, Taylor JA, Kim DW, Place ES, Rogers TF, Mank JE, Cooney CR, Wright AE. 2022. Detecting signatures of selection on gene expression. Nat Ecol Evol 6:1035–1045.

Pšeničnik A, Slemc L, Avbelj M, Tome M, Šala M, Herron P, Shmatkov M, Petek M, Baebler Š, Mrak P, et al. 2024. Oxytetracycline hyper-production through targeted genome reduction of *Streptomyces rimosus*. mSystems 9:e00250–24.

Rowlands RT. 1984. Industrial strain improvement: rational screens and genetic recombination techniques. Enzyme Microb Tech 6:290–300.

Santamarta I, Rodriguez-Garcia A, Perez-Redondo R, Martin J, Liras P. 2002. CcaR Is an Autoregulatory Protein That Binds to the ccaR and cefD-cmcI Promoters of the Cephamycin C-Clavulanic Acid Cluster in *Streptomyces clavuligerus*. Journal of Bacteriology 184.

Saxer G, Doebeli M, Travisano M. 2010. The Repeatability of Adaptive Radiation During Long-Term Experimental Evolution of *Escherichia coli* in a Multiple Nutrient Environment. PLoS ONE 5: e14184.

Schniete JK, Cruz-Morales P, Selem-Mojica N, Fernández-Martínez LT, Hunter IS, Barona-Gómez F, Hoskisson PA. 2018. Expanding Primary Metabolism Helps Generate the Metabolic Robustness To Facilitate Antibiotic Biosynthesis in *Streptomyces*. Mbio 9: e02283–17.

Seemann T. 2014. Prokka: rapid prokaryotic genome annotation. Bioinformatics 30:2068–2069.

Song JY, Jeong H, Yu DS, Fischbach MA, Park H-S, Kim JJ, Seo J-S, Jensen SE, Oh TK, Lee KJ, et al. 2010. Draft Genome Sequence of *Streptomyces clavuligerus* NRRL 3585, a Producer of Diverse Secondary Metabolites. J Bacteriol 192:6317–6318.

Steensels J, Gallone B, Voordeckers K, Verstrepen KJ. 2019. Domestication of Industrial Microbes. Curr Biol 29: R381–R393.

Stoltzfus A. 2021. Mutation, Randomness, and Evolution. Oxford University Press.

Tahlan K, Park HU, Wong A, Beatty PH, Jensen SE. 2004. Two Sets of Paralogous Genes Encode the Enzymes Involved in the Early Stages of Clavulanic Acid and Clavam Metabolite Biosynthesis in *Streptomyces clavuligerus*. Antimicrob. Agents Chemother. 48:930–939.

Tidjani A-R, Lorenzi J-N, Toussaint M, Dijk E van, Naquin D, Lespinet O, Bontemps C, Leblond P. 2019. Massive Gene Flux Drives Genome Diversity between Sympatric *Streptomyces* Conspecifics. mBio 10:e01533–19.

Uchida T, Miyawaki M, Kinashi H. 2003. Chromosomal Arm Replacement in *Streptomyces griseus*. J. Bacteriol. 185:1120–1124.

Ünsaldı E, Kurt-Kızıldoğan A, Voigt B, Becher D, Özcengiz G. 2017. Proteome-wide alterations in an industrial clavulanic acid producing strain of *Streptomyces clavuligerus*. Synthetic Syst Biotechnology 2:39–48.

Wagner A. 2023. Evolvability-enhancing mutations in the fitness landscapes of an RNA and a protein. Nat. Commun. 14:3624.

Wilson DJ, Consortium TCr, Crook DW, Peto TEA, Walker AS, Hoosdally SJ, Cruz ALG, Carter J, Grazian C, Earle SG, et al. 2020. GenomegaMap: within-species genome-wide dN/dS estimation from over 10,000 genomes. Mol Biol Evol. 37:2450–2460.

Wu X, Roy KL. 1993. Complete nucleotide sequence of a linear plasmid from *Streptomyces clavuligerus* and characterization of its RNA transcripts. J Bacteriol 175:37–52.

Wu W, Leblanc SKD, Piktel J, Jensen SE, Roy KL. 2006. Prediction and functional analysis of the replication origin of the linear plasmid pSCL2 in *Streptomyces clavuligerus*. Can J Microbiol. 52:293–300.

Yanai K, Murakami T, Bibb M. 2006. Amplification of the entire kanamycin biosynthetic gene cluster during empirical strain improvement of *Streptomyces kanamyceticus*. Proceedings of the National Academy of Sciences of the United States of America 103:9661–9666.

Zhu M, Dai X. 2024. Shaping of microbial phenotypes by trade-offs. Nat. Commun. 15:4238.

